# Saturated lipid stress attenuates mitochondrial genome synthesis in human cells

**DOI:** 10.1101/2025.07.28.667051

**Authors:** Casadora Boone, Sophie Judge, Ahmad Shami, Bezawit Danna, Andrea B. Ball, Tejashree Pradip Waingankar, Hera Saqub, Ajit S. Divakaruni, Samantha C. Lewis

## Abstract

Fatty acids are trafficked between organelles to support membrane biogenesis and act as signaling molecules to rewire cellular metabolism in response to starvation, overnutrition, and environmental cues. Mitochondria are key cellular energy converters that harbor their own multi-copy genome critical to metabolic control. In homeostasis, mitochondrial DNA (mtDNA) synthesis is coupled to mitochondrial membrane expansion and division at sites of contact with the endoplasmic reticulum (ER). Here, we provide evidence from cultured hepatocytes that mtDNA synthesis and lipid droplet biogenesis occur at spatially and functionally distinct ER-mitochondria membrane contact sites. We find that, during saturated lipid stress, cells pause mtDNA synthesis and mitochondrial network expansion secondary to rerouted fatty acid trafficking through the ER and lipid droplet biogenesis, coincident with a defect in soluble protein import to the ER lumen. The relative composition of fatty acid pools available to cells is critical, as monounsaturated fatty acid supplementation rescued both ER proteostasis and mtDNA synthesis, even in the presence of excess saturated fat. We propose that shutoff of mtDNA synthesis conserves mtDNA-to-mitochondrial network scaling until cells can regain ER homeostasis.

**Summary:** Overnutrition of cultured human cells causes endoplasmic reticulum dysfunction, which downregulates mitobiogenesis in turn by constraining mtDNA synthesis.

## Introduction

Mitochondria play a central role in lipid metabolism through beta-oxidation of fatty acids for ATP production, as well as through interactions with other lipid metabolizing organelles including the endoplasmic reticulum, lipid droplets, peroxisomes, and lysosomes [1–3]. Lipid droplets (LDs) are the major sites of neutral lipid storage in cells. Membrane contact sites between mitochondria, LDs, and the endoplasmic reticulum (ER) define a subset of peri-droplet mitochondria that preferentially participate in ATP production via fatty acid beta-oxidation [4–7]. Cellular nutrient and energy demand modulate fatty acid flux into and out of the LD compartment and thus the balance between lipid storage and usage, as well as the abundance of peri-droplet mitochondria [2, 8–9].

Beyond energy conversion, fatty acids are trafficked between organelles to support membrane biogenesis and act as signaling molecules that are needed to rewire cellular metabolism in starvation, overnutrition, and in response to environmental cues [1, 10–17]. These signaling events, and subsequent execution of metabolic programs, depend on membrane contact sites between mitochondria, LDs, and the ER. Disruption of lipid homeostasis, particularly excess cellular uptake of saturated fatty acids, remodels lipid trafficking through LDs and ER, impinging organellar biogenesis and protein quality control functions [18–20]. This lipotoxic insult activates cellular stress responses that monitor cytoplasmic protein quality control, including the ER unfolded protein response (ER-UPR) and the broader integrated stress response (ISR) [10, 21–24].

Lipotoxicity also leads to mitochondrial dysfunction, although the mechanisms by which saturated lipid stress influences mitobiogenesis are not clear [7–8; 25–26]. Mitochondria harbor their own genome, mitochondrial DNA (mtDNA) which encodes essential subunits of the electron transport chain complexes I, III, IV, and V [27]. MtDNA is maintained by dedicated proteins that constitute the mitochondrial replisome, minimally consisting of a DNA polymerase (POLG1 and POLG2), single-stranded binding protein (SSBP1), and helicase (TWINKLE) [27–29]. In humans, deleterious mtDNA mutations are linked to increased lipid deposits in the skeletal muscle and liver [25, 30]. In mice, mtDNA replication defects have been associated with an overall decrease in lipid production that correlates with peripheral neuropathy [31]. Moreover, both saturated fatty acid treatment in cell culture and obesity in mice are associated with an increased load of mtDNA lesions and mitochondrial membrane herniation, leading to the release of pro-inflammatory mtDNA into the cytosol [32–38]. This suggests that mtDNA synthesis and integrity are coupled to lipid metabolism by largely unknown crosstalk machineries.

Lipids move between ER, LDs, and mitochondria via membrane contact sites (MCS) that serve as platforms for lipid and ion transfer, lipid droplet biogenesis, mitochondrial fission, mtDNA synthesis, and mtDNA nucleoid partitioning in proliferating cells [39–46]. The abundance and size of ER-mitochondria MCS are responsive to nutrient availability, ISR activation, as well as the progression of mtDNA synthesis [47–52]. Whether these disparate functions occur simultaneously at all ER-mitochondria MCS, or are distributed amongst spatially or regulated classes of such, is not well understood.

Here we investigate how two well-characterized MCS functions, lipid droplet biogenesis and mtDNA synthesis, are spatially organized in cells and the consequences of organellar reorganization during lipid stress. We find that the subset of mitochondria engaged in mtDNA synthesis are distinct from those participating in tripartite interactions for LD biogenesis. During stress induced by excess saturated fatty acid, recruitment of the mtDNA polymerase subunit POLG2 to mtDNA is impaired and mtDNA synthesis is attenuated, concurrent with ER dilation. ER morphology, POLG2 recruitment, and mtDNA synthesis are rescued by rebalancing fatty acid pools via supplementation with a monounsaturated fatty acid. We propose that endomembrane remodeling in lipid stress promotes enhanced lipid neutralization and storage at the cost of mtDNA synthesis; thus, prioritizing metabolic adaptation over mitochondrial biogenesis.

## Results

### MtDNA synthesis and LD maintenance occur at distinct ER-mitochondria contacts

To identify the positioning of lipid droplets relative to sites of mtDNA synthesis at ER-mitochondria contacts, we transiently transfected human hepatoma (Huh7) cells labeled with BODIPY-665/676 with three fluorescent reporter proteins: the ER marker KDEL-mRuby, mitochondrial matrix-targeted BFP, and tagged mtDNA polymerase subunit POLG2-GFP. Airyscan fluorescence microscopy revealed POLG2-GFP foci in a subset of mitochondria, that were enriched at points of ER-mitochondria colocalization, as previously shown in other immortalized mammalian cell lines (**Figure 1A, top right**) [43]. On average, each cell contained 120 POLG2-GFP foci that were sparsely distributed among the mitochondrial population, where they often colocalized with, or were adjacent to, ER tubules engaged in mitochondrial constriction (**Figure 1A-B**). We observed Bodipy-labeled LDs at tripartite intersections with ER and a subset of mitochondria throughout the cytoplasm (**Figure 1A, lower right**). To determine whether sites of contact with LDs were spatially linked to sites of mtDNA synthesis, we segmented regions of colocalization between ER and mitochondria marked by POLG2-GFP and quantified the number that also intersected with LDs (**Figure 1C**). Fewer than 20% of ER-mitochondria contacts were associated also with LDs, indicating that ER-mitochondria-LD tripartite contacts are distinct from sites of mtDNA replication.

**Figure 1.**
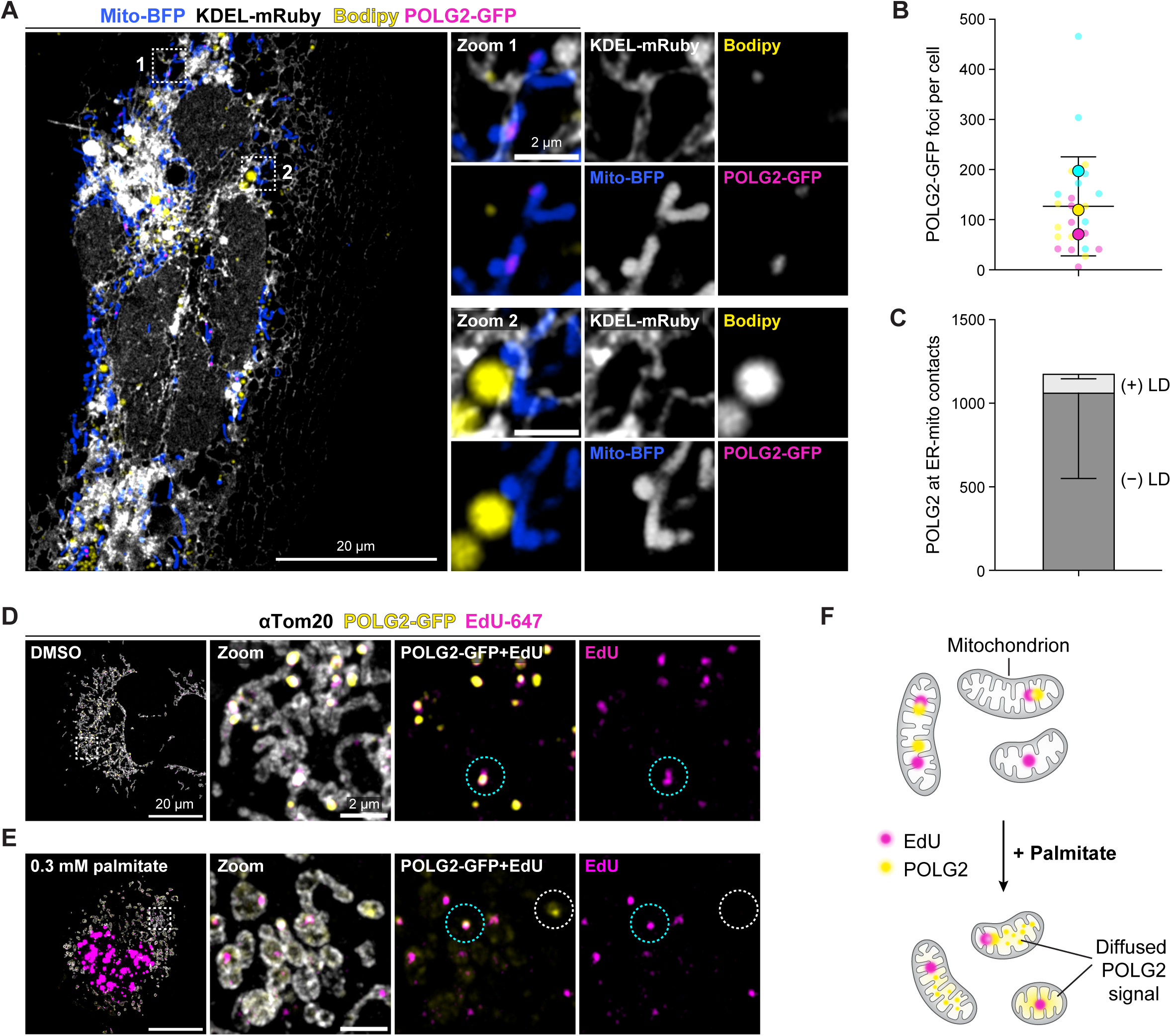
Mitochondrial genome synthesis and lipid droplet maintenance occur at distinct ER-mitochondria contacts. **(A)** Huh-7 cell transiently transfected with markers KDEL-mRuby (gray), mito-BFP (blue), POLG2-GFP (magenta), and stained with the far red neutral lipid staining dye BODIPY 665/676 (yellow). Scale bar 20 μm. Zoom 1, ER-mitochondria colocalization at replicating mtDNA nucleoid. Zoom 2, Tripartite contact between ER-mitochondria-lipid droplet. Scale bar 2 μm. **(B)** Quantification of POLG2-GFP foci per cell, N =3,167 foci from 25 cells across 3 replicates. **(C)** Quantification of the number of POLG2-GFP positive ER-mitochondrial intersections with or without lipid droplets association. N=3,529 intersections from 25 cells across 3 replicates. **(D-E)** Fluorescent labeling of nascent DNA in mitochondria of Huh-7 cells (EdU-AlexaFluor647, magenta) transiently transfected with POLG2-GFP (yellow), and immunolabeled with anti-Tom20 (gray), then treated for 24 hours with DMSO (D) or 0.3 mM PA (E). Cyan circles indicate colocalization between POLG2-GFP and EdU; white circles denote GFP signal without EdU-AlexaFluor647. N = 37 cells (DMSO), or 34 cells (0.3 mM PA) from 3 biological replicates. Scale bars, 10 μm and 2 μm. **(F)** Schematic comparison of mitochondrial EdU and POLG2-GFP signal in control versus PA-treated cells.

To validate these observations we employed a genetically-encoded reporter for LD biogenesis, GFP-tagged Seipin. Seipin forms homo-oligomeric complexes that support ER-LD contact site maintenance and the trafficking of neutral, esterified lipids into nascent LDs [4; 51]. We marked active mtDNA nucleoids in the same cells using an mKate-tagged allele of SSBP1, the mitochondria-specific single-stranded binding protein required for mtDNA synthesis and transcription (**Figure S1**). Seipin-GFP formed fluorescent rings around a subset of spherical, perinuclear LDs, most of which were distal to mitochondria marked by SSBP1-mKate. These results are consistent with a model in which subcellular compartmentalization maintains a subpopulation of mitochondria engaged in mtDNA synthesis distinct from another subpopulation associated with LDs in Huh7 cells.

Given that both mitochondria and LDs make functionally essential contacts with the ER, we considered that ER remodeling to support the biogenesis of one organelle interaction partner could involve a reciprocally altered interaction with the other. The lipid composition of mitochondrial membranes informs mitochondrial division by the mitochondrial division dynamin Drp1 via its receptors, tuning mitochondrial fission [43, 44]. Mitochondrial morphology and fatty acid oxidation (FAO) rates are intimately linked, with greater FAO leading to fragmentation of the mitochondrial network [53]. Moreover, a high fat diet entrains mtDNA depletion, suggesting that mtDNA maintenance is functionally linked to lipid homeostasis across different scales from organelles, to cells, to tissues and the whole organism [Reviewed in 25]. To address this hypothesis, we tested whether saturated lipid stress induced via excess of the saturated fatty acid palmitate (PA), would impact mtDNA synthesis at ER-mitochondria contacts. PA is a long-chain saturated fatty acid linked to perturbed mtDNA integrity in humans with metabolic disease, as well as to liver lipotoxicity, steatosis, and systemic inflammation [19, 38, 54–55].

We pulse-labeled cells expressing POLG2-GFP with the synthetic thymidine analog 5-ethynyl-2-deoxyuridine (EdU), then fixed and stained them to detect EdU-AlexaFluor647, the endogenous mitochondrial outer membrane protein Tom20, or GFP, by immunofluorescence (**Figure 1D-G**). In cells treated with only the DMSO vehicle, distinct POLG2-GFP puncta colocalized with EdU foci in tubular mitochondrial networks consistent with the phenotype of the marker in live cells (**Figure 1D**). In contrast, POLG2-GFP was diffusely localized within mitochondria in cells cultured in PA-supplemented medium, (**Figure 1E**) in which only a minority of POLG2-GFP puncta colocalized precisely with EdU (**Figure 1E, cyan circle**). When POLG2 puncta were present, they often did not colocalize with EdU (**Figure 1E, white circle**). In parallel, mitochondria were fragmented upon exposure to this nutrient excess, consistent with previous reports [56–58]. These observations indicate that the recruitment of at least one component of the mtDNA replisome to mtDNA at ER-mitochondria contacts is perturbed in cells fed excess PA (**Figure 1F**). Thus, we sought to quantify mitochondrial function, mtDNA copy number and mtDNA synthesis systematically in our lipid stress model.

### Lipid accumulation model to probe effects on mitochondrial function

We established a lipotoxicity model in which to examine reporters of organelle morphometrics, mtDNA synthesis, and fatty acid trafficking in cells subject to an accumulation of excess PA. Palmitate cannot cross mitochondrial membranes directly, but must be trans-esterified to a neutral lipid prior to mobilization into droplets or mitochondrial uptake via the carnitine palmitoyltransferase (CPT1) shuttle [59]. Huh7 cells cultured in 0.1-0.3 mM PA for 24 h exhibited reduced Mitotracker fluorescence intensity consistent with impaired mitochondrial inner membrane potential and accumulated LDs, as shown by Bodipy 493/503 staining (**Figure 2A-C**). Mitochondrial network size was also reduced in PA-treated cells relative to non-treated and DMSO vehicle-treated controls (**Figure 2D**). We validated the reduction in membrane potential by quantifying the fluorescence intensity of tetramethyl rhodamine ester (TMRE), which was also significantly reduced (**Figure 2E-F**).

**Figure 2.**
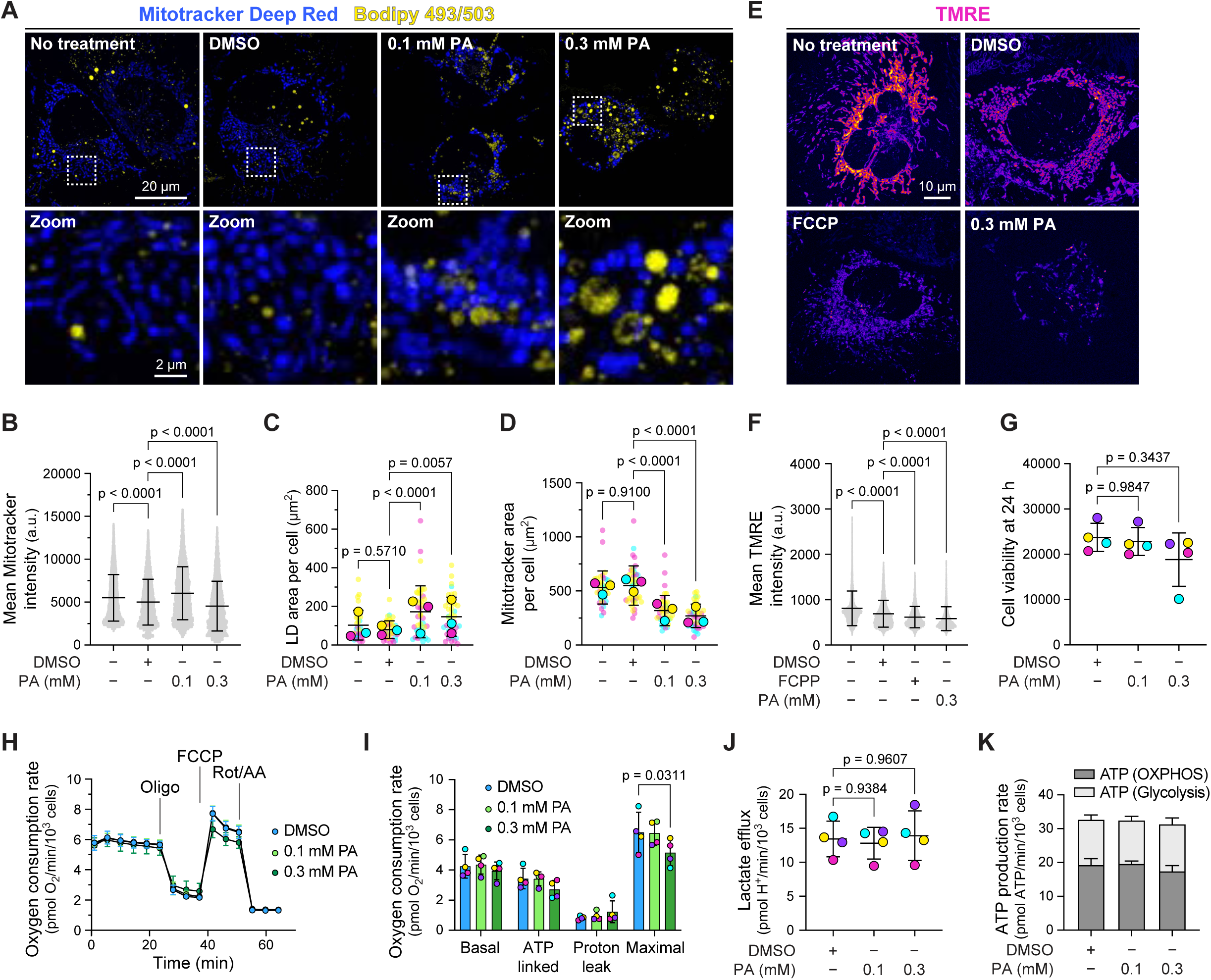
Lipid accumulation model to probe effects on mitochondrial function. **(A)** Untreated Huh-7 cells versus DMSO vehicle, or palmitate and stained with BODIPY 493/503 (yellow), and MitoTracker Deep Red (blue). Scale bars, 10 μm. **(B)** Mean Mitotracker Deep Red fluorescence intensity in segmented mitochondria, in the presence or absence of palmitate, quantified from cells in (A). N = 54 cells (NT), 52 cells (DMSO), 64 cells (0.1 mM PA), 47 cells (0.3 mM PA) cells, from 3 biological replicates. **(C)** Lipid droplet area (μm^2^) per cell, quantified from cells in (A). One-way ANOVA (Dunnett’s multiple comparisons test). **(D)** Segmented mitochondrial area (μm^2^) per cell, quantified from cells in (A). **(E)** Untreated Huh-7 cells versus DMSO vehicle, FCCP positive control, or palmitate and stained with TMRE (fire LUT) and BODIPY 493/503. Scale bar, 10 μm. **(F)** Mean TMRE fluorescence intensity in segmented mitochondria, quantified from cells in (E). N = 34 cells (NT), 32 cells (DMSO), 25 cells (FCCP), and 26 cells (0.3 mM PA), from 3 biological replicates. **(G)** Mean ± SD of cell count in DMSO versus PA after 24 h. N = 94976 (DMSO), 91345 (0.1 mM PA), 75341 (0.3 mM PA) cells from 4 biological replicates. **(H)** Representative respirometry trace of cells treated as in (G), N = 4 biological replicates. **(I)** Aggregate Basal, ATP-linked, Proton leak, and Maximal respiration in intact Huh7, as in (G), N = 4 biological replicates. **(J)** Lactate efflux rate from respirometry experiments in (H-I), one-way ANOVA (Dunnett’s multiple comparisons test). **(K)** ATP production rates calculated for Huh7 cells, N = 4 biological replicates.

Despite these changes, cell viability remained high, indicating that our treatment regime was sufficient to influence cell metabolism without crossing a threshold of dysfunction that would instigate a commitment to cell death (**Figure 2G**). Cells cultured in PA exhibited small yet statistically significant changes in the oxygen consumption rate (**Figure 2H-I**) but not in the proportion of ATP generated by oxidative phosphorylation versus glycolysis, or lactate efflux (**Figure 2J-K**). Thus, PA treatment led to an expansion of cellular lipid stores accompanied by measurable impacts on mitochondrial network organization without confounding effects from bioenergetic dysfunction.

### Saturated lipid stress attenuates mtDNA synthesis

We next sought to determine the impact of cell culture in the presence of excess saturated fatty acid on mtDNA homeostasis using direct metabolic labeling of DNA synthesis. Given the altered POLG2 recruitment to nucleoids in PA-treated cells, we considered that mtDNA might be depleted from mitochondrial networks due to decreased replication efficiency. Alternatively we hypothesized that, if overall mitochondrial network growth were abrogated, suppression of mtDNA synthesis could serve to maintain the scaling of mitochondrial genome copies to network volume until the lipid stress were relieved.

To test our hypothesis, we quantified the number of mitochondrial DNA and mitochondrial EdU foci in cells cultured in the presence of pharmacologic inhibitors of diacylglycerol acetyltransferase enzymes DGAT1 and DGAT2 for 48 h, with or without the 24 hour 0.3 mM PA treatment. Once long-chain saturated fatty acids enter cells, they are esterified in the ER then accumulate as neutral lipids in nascent LDs that then lens and bud from the ER membrane bilayer into the cytosol. LD lensing and budding are stabilized by DGAT1 and DGAT2, respectively. If the efficient neutralization and storage of free fatty acids were a requirement for mtDNA replication, then DGAT1/DGAT2 inhibition should exacerbate the effect of PA on mtDNA synthesis.

We found that acute PA exposure alone had no effect on mtDNA/dsDNA foci or their density in mitochondrial networks normalized for network size (**Figure 3A-B**), which was validated by quantitative PCR of mtDNA copy number per cell (**Figure S2A**). In contrast, we observed a significant simultaneous decrease in the proportion of mitochondrial EdU foci in the same cells (**Figure 3C**). Strikingly, the density of EdU foci, but not dsDNA foci, in mitochondrial networks was significantly decreased, indicating that during saturated lipid stress cells maintain their mtDNA pool in a quiescent state (**Figure 3D**).

**Figure 3.**
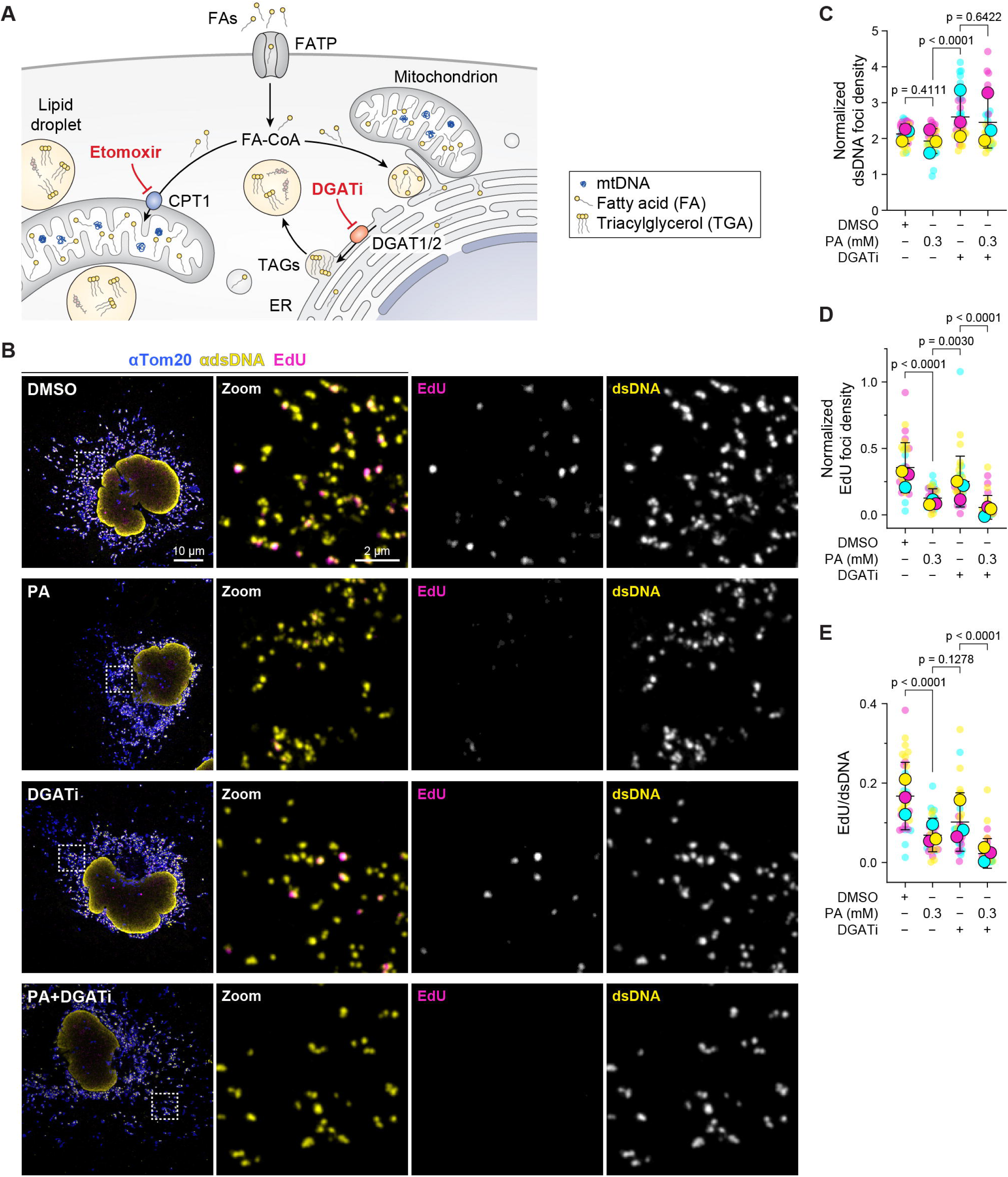
Saturated lipid stress attenuates mtDNA synthesis. **(A)** Schematic of fatty acid trafficking in cells upon treatment with inhibitors of DGAT1/2 (DGATi) or mitochondrial fatty acid uptake by CPT1 (Etomoxir). **(B)** Huh-7 cells treated with DMSO vehicle, T863/PF-06424439 (DGATi), 0.3 mM PA or both, then pulse-labeled with EdU-AlexaFluor647 (magenta) and immunolabeled with anti-Tom20 (blue) and anti-dsDNA (yellow). Scale bar, 10 μm. **(C)** Quantification of mitochondrial dsDNA foci per cell, normalized to segmented mitochondrial area. **(D)** Quantification of mitochondrial EdU-647 foci per cell, normalized to segmented mitochondrial area. **(E)** Calculation of the proportion of mtDNA nucleoids engaged in DNA synthesis on a per-cell basis, from (C-D). One-way ANOVA (Dunnett’s multiple comparisons test). Cyan, magenta and yellow data points represent individual cells from each replicate. For (C-E), N=32 cells (NT), 34 cells (DMSO), 36 cells (0.3 mM PA), 36 cells (DGATi), and 36 cells (DGATi + 0.3 mM PA), from 3 biological replicates.

In the same experiment we used pharmacological manipulation of fatty acid trafficking between ER, LDs, and mitochondria to determine which FA routes were relevant to EdU labeling of mtDNA in our model. Combinatorial DGAT1 and DGAT2 inhibition was sufficient to reduce mitochondrial DNA synthesis, and further exacerbated the attenuation induced by PA treatment alone (**Figure 3C**), demonstrating that inhibition of TAG biosynthesis is sufficient to perturb mtDNA.

These effects were concurrent with a significant reduction in LDs per cell, demonstrating the effectiveness of DGAT1/2 inhibition (**Figure S2B-C**). In contrast, direct inhibition of mitochondrial fatty acid uptake using etomoxir, a small molecule inhibitor of CPT1, had no impact on mitochondrial dsDNA foci abundance or EdU labeling (**Figure S2D-F**). These results demonstrate that mtDNA synthesis is not co-regulated with mitochondrial fatty acid uptake, while suggesting that the accumulation of free fatty acids in the ER when they can’t be trafficked into LDs is more relevant. These findings led us to wonder whether lipid accumulation in the ER may impinge on mtDNA synthesis via altered ER function and/or MCS with mitochondria.

### Decreased mtDNA synthesis is correlated with rerouted FA trafficking through the ER

High fat diet causes ER dilation and MCS remodeling in mouse models as well as human liver tissue [20, 47, 50]. Moreover, recent work indicates that the quantity of stable ER-mitochondria contacts in cultured cells is tuned to the level of integrated stress response (ISR) activation during the ER unfolded protein stress response (ER-UPR) [60]. Thus, we asked whether excess saturated fatty acids impinge on ER proteostasis.

We first assessed the abundance of ER stress marker proteins CHOP, XBP1s, and GRP78 after 24 h cell culture in PA by immunoblot (**Figure 4A**). We observed weak induction of XBP1s and CHOP in cells supplemented with 0.3 mM PA, yet no change in GRP78 protein levels, consistent with mild ER-UPR activation. We next assessed ER protein import competence by comparing the localization of fluorescent reporter constructs targeted to the ER membrane versus the ER lumen via transient transfection with GFP-SEC61b and KDEL-mRuby, respectively. We predicted that ER dysfunction would result in reduced import of soluble proteins to the ER lumen, specifically. Consistently, GFP-Sec61b and KDEL-mRuby were robustly colocalized in control cells (Pearson’s R = 0.90), yet KDEL-mRuby fluorescence was largely cytoplasmic in cells cultured in excess palmitate (Pearson’s R = 0.13) (**Figure 4B**). These data validate that saturated lipid stress compromises ER function.

**Figure 4.**
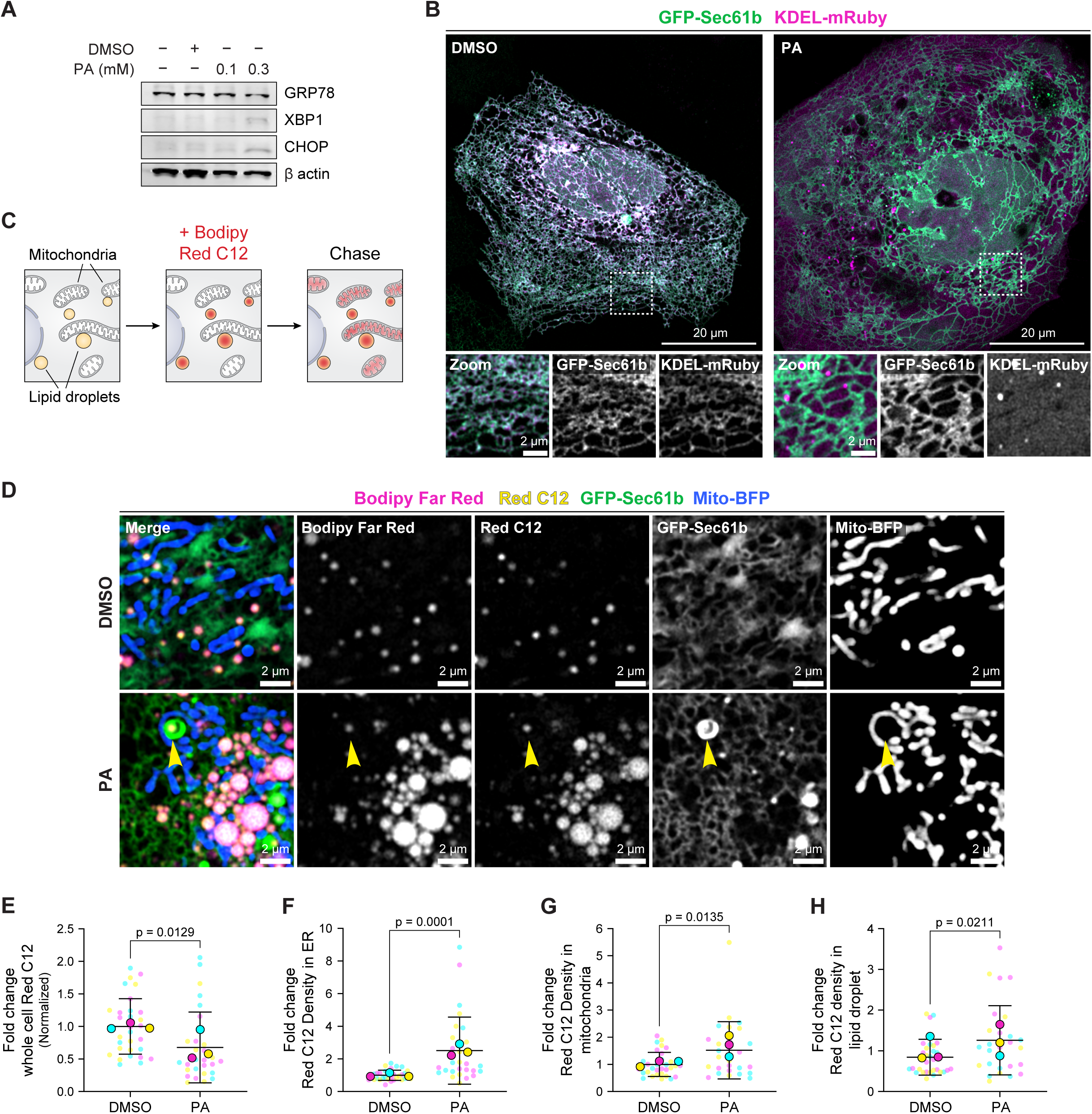
Impaired mtDNA synthesis is correlated with rerouted FA trafficking. **(A)** Untreated Huh7 cells versus DMSO or PA, analyzed by immunoblot for ER stress marker protein abundance relative to beta-actin. **(B)** Huh-7 cells transiently transfected with KDEL-mRuby and GFP-Sec61b, treated with DMSO (left) or palmitate (right). Scale bars, 20 μm; 2 μm. **(C)** Schematic of BODIPY 558/568 (Red C12) loading into cells in culture. **(D)** Huh-7 cells transiently transfected with Mito-BFP and GFP-Sec61b, labeled with Bodipy Far Red and Red C12, treated with DMSO (top row) or palmitate (bottom row). Arrow indicates Colocalized Bodipy Far Red and Red C12 within a bulb of ER membrane. Scale bar, 2 μm. **(E)** Quantification of total Red C12 fluorescence per cell/uptake in cells from panel D. N = 32 cells (DMSO), 28 cells (PA) from 3 replicates. Quantification of Red C12 fluorescence density within segmented ER **(F)**, mitochondria **(G)**, and lipid droplets **(H)** in cells labeled as in D. Significance determined via two-tailed unpaired t-test.

To monitor the intracellular fates of exogenous lipids and how they may be rerouted during nutrient excess, we labeled cells with the fluorescent fatty acid analogue Bodipy 558/568 C12 (Red C12) in complete media with or without PA supplementation (**Figure 4C**). In control cells transiently transfected with GFP-Sec61b and mito-BFP, and labeled with Bodipy Far Red, Red C12 colocalized with punctate LDs (**Figure 4D**). In contrast, PA-treated cells exhibited a greater proportion of cellular Red C12 fluorescence colocalized with ER and mitochondria. Concurrent with Red C12 accumulation in non-LD organelles, we noted Red C12- and Bodipy Far Red-enriched domains enclosed by whorls of ER membrane in contact with mitochondria within PA-treated cells (**Figure 4D, yellow arrow**), suggestive of nascent LDs trapped in the ER. Cellular uptake of Red C12 was consistent across replicates (**Figure 4E**), and thus we quantified the density of Red C12 fluorescence in each organelle upon culture in DMSO versus PA (**Figure 4F-H**). These analyzes revealed a 3-fold increase in Red C12 density colocalized with ER upon culture in palmitate, whereas mitochondrial and LD Red C12 density was increased 1.1 to 1.5-fold. Consistent with the impaired mtDNA synthesis in DGAT1/2i cells, these results indicate that impaired fatty acid transfer from ER to LD is likely to contribute to mitochondrial dysfunction as lipids accumulate in ER and mitochondria.

### Rebalanced fatty acid ratios rescue mtDNA synthesis

We next considered whether ER function and attenuated mtDNA synthesis could be rescued by rebalancing the ratio of saturated to unsaturated fatty acids available to cells. The monounsaturated fatty acid oleic acid (OA) exerts cytoprotective effects against PA-induced cell death and ER stress in multiple mammalian hepatocarcinoma cell lines and in skeletal muscle cells via increased triacylglycerol (TAG) flux and mitochondrial beta-oxidation [55; 61; 62]. Thus, we asked whether supplementation of cell culture medium with 0.3 mM OA was sufficient to rescue ER lumenal protein import, mtDNA synthesis, or both. We reasoned that, if mtDNA synthesis is attenuated downstream of ER dysfunction, rescue of ER protein import by addition of oleic acid should also rescue mtDNA replication.

We transiently transfected Huh7 cells with KDEL-mRuby, mitochondrial matrix-targeted BFP, and POLG2-GFP then incubated them with PA, OA, or both simultaneously. As we had previously observed, mitochondrial POLG2-GFP signals were diffuse in cells treated with 0.3 mM PA alone, suggestive of decreased recruitment of the replisome machinery to mtDNA nucleoids (**Figure 5A**). Treatment with OA alone had no discernible impact on POLG2-GFP foci as compared to no-treatment or DMSO controls. In contrast, distinct POLG2-GFP puncta were present in cells treated with PA and OA simultaneously, despite the fact that mitochondria remained fragmented in some cells (**Figure 5A, Figure S3**). Strikingly, the KDEL-mRuby import to the ER was completely rescued by simultaneous culture in OA and PA (**Figure 5A, see lower left panel**).

**Figure 5.**
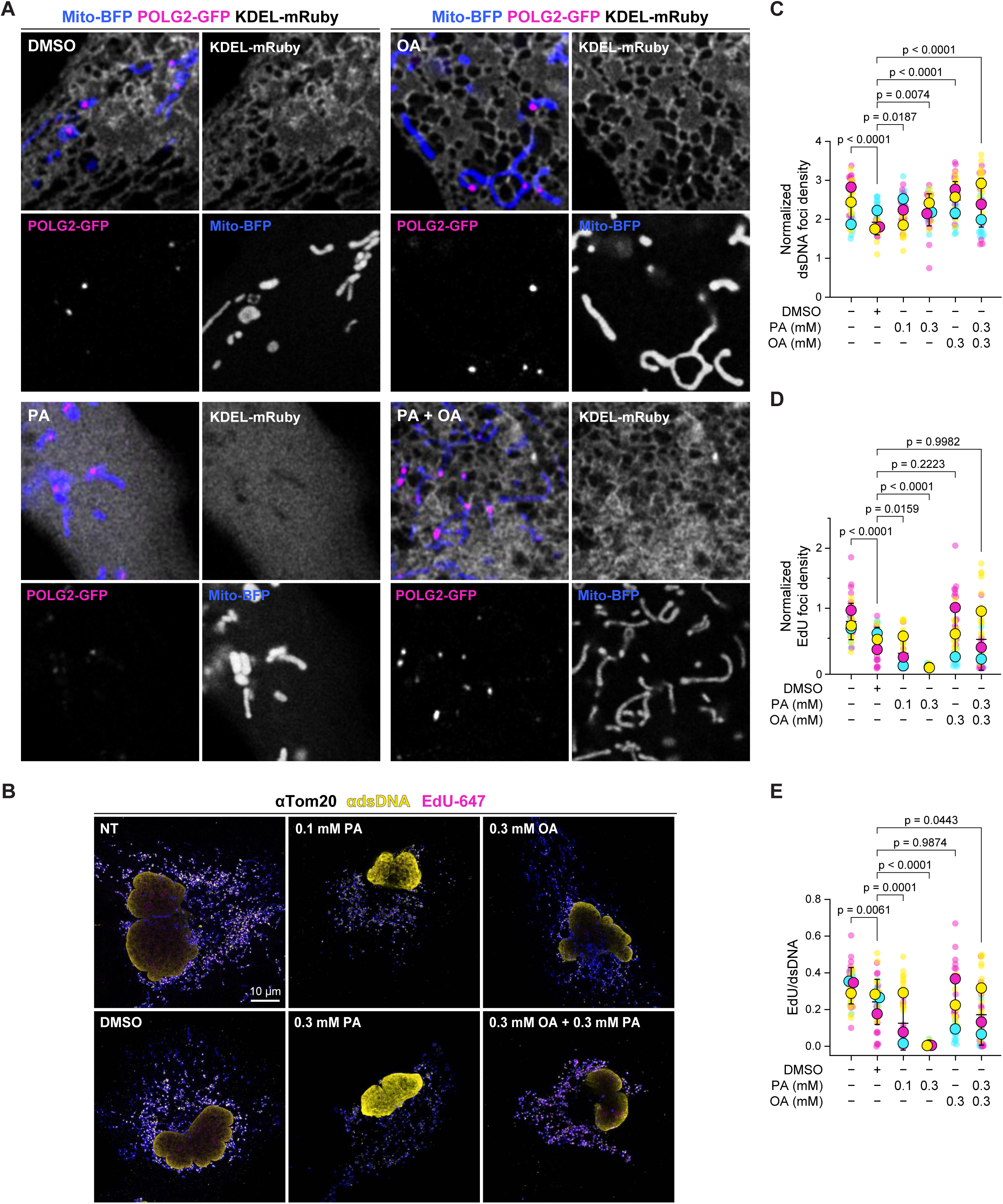
Rebalanced fatty acid ratios rescue mtDNA synthesis. **(A)** Huh-7 cells transiently transfected with Mito-BFP, POLG2-GFP, and KDEL-mRuby and treated with DMSO, oleic acid (OA), palmitate (PA), or both (OA+PA). **(B)** Huh-7 cells treated with DMSO vehicle, OA, PA, or both, then pulse-labeled with EdU-AlexaFluor647 (magenta) and immunolabeled with anti-Tom20 (blue) and anti-dsDNA (yellow). Scale bar, 10 μm. **(C)** Quantification of mitochondrial dsDNA foci per cell normalized to segmented mitochondrial area reported by Tom20 (μm^2^), data from (B). **(D)** Quantification of mitochondrial EdU-647 foci per cell normalized to segmented mitochondrial area, data from (B). **(E)** Calculation of the proportion of mtDNA nucleoids engaged in DNA synthesis on a per-cell basis, from (B), one-way ANOVA (Dunnett’s multiple comparisons test). Cyan, magenta and yellow data points represent individual cells from each replicate. For (C-E), N = 50 cells (NT), 55 cells (DMSO), 61 cells (0.1 mM PA), 52 cells (0.3 mM PA), 52 cells (0.3 mM OA), 49 cells (0.3 mM OA+PA), from 3 biological replicates.

**Figure 6.**
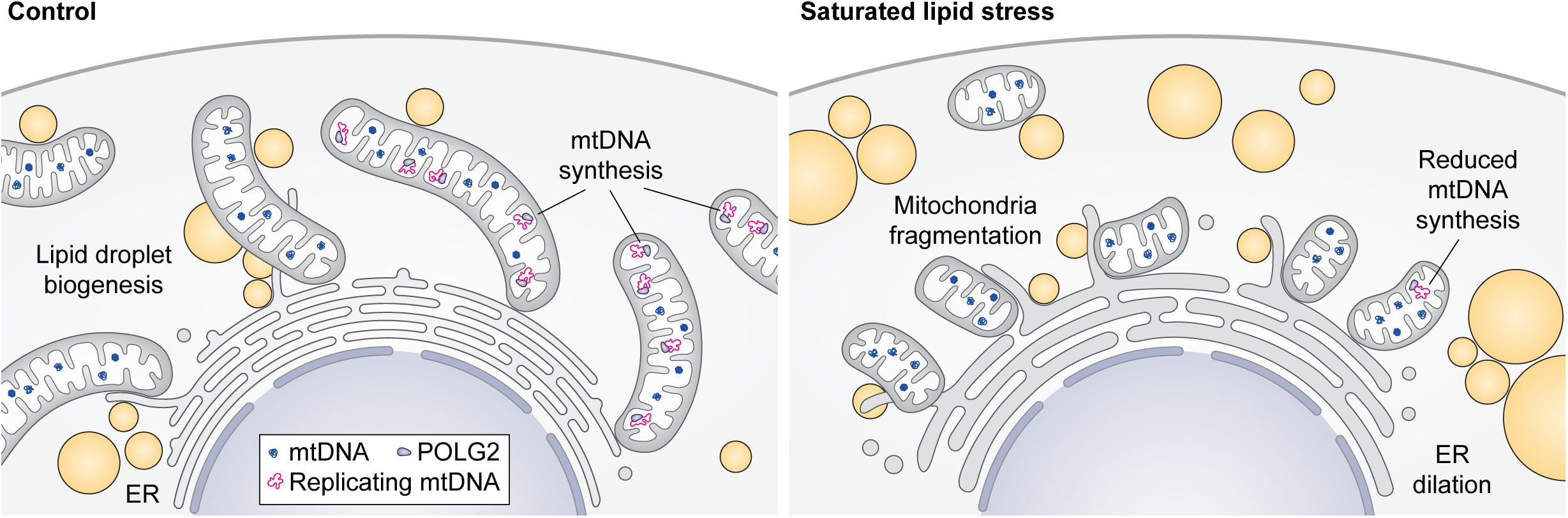
Saturated lipid stress attenuates mtDNA synthesis downstream of ER dysfunction.

Consistent with our previous observation of the minor effects of PA supplementation on mitochondrial respiration and cell viability, analysis of ER and mitochondrial mass by western blot detection of endogenous Calreticulin and Tom20 revealed no differences between cells treated with DMSO, PA, OA, or OA and PA simultaneously (**Figure S4A**), indicating that KDEL import was rescued without significant change in ER mass per cell. Further, mitochondrial network size (**Figure S4B**) and mtDNA synthesis (**Figure S4C-E**) were unaffected by inhibition of the ISR using the potent small molecule ISRIB, either alone or in combination with PA treatment. ISRIB blocks eIF2a-dependent adaptation to the ER-UPR, thus, these data indicate that our observations are largely independent of the suppression or activation of cytosolic protein translation in stress [60]. Taken together, these results are consistent with the hypothesis that ER dysfunction during lipotoxic insult disrupts interactions with mitochondria and, as a consequence, mtDNA synthesis.

Finally, we validated that rebalancing saturated and unsaturated fatty acids in the cell culture medium rescued mtDNA labeling by EdU. We quantified the number of mitochondrial double-stranded DNA foci and EdU foci per cell in samples cultured in PA, OA, or both. Cell culture in excess OA alone had no impact on the number of mitochondrial dsDNA foci per cell or EdU labeling; the density of replicating mtDNAs in mitochondria was also unchanged (**Figure 5B-D**). However, concomitant supplementation with excess PA and OA rescued EdU labeling of mtDNA (**Figure 5D**). The proportion of mtDNA nucleoids engaged in replication was not significantly different in OA+PA-treated cells from untreated or DMSO-treated cells (**Figure 5E, Figure S5**).

In sum, these findings indicate that the relative composition of the fatty acid pools available to cells rather than their absolute amounts influences the proportion of mtDNA nucleoids engaged in DNA synthesis, which is determined by the overall function of the ER and its interplay with mitochondria in response to saturated lipid stress.

## Discussion

Cell homeostasis requires coordination between multiple processes in response to intrinsic and extrinsic cues. Given the spatial distribution of biochemical functions into distinct organelles, MCSs between them serve as hubs for such coordination. Here, we provide evidence that mtDNA synthesis and LD biogenesis occur at spatially and functionally distinct intersections with the ER, with potential implications for understanding pathological lipotoxicity.

We posit that peri-droplet mitochondria are a distinct subpopulation excluded from mtDNA synthesis and mitochondrial biogenesis. Peri-droplet mitochondria are well-characterized players in FA trafficking and are critical to cells successfully making the metabolic shift to beta-oxidation when substrates suitable for breakdown by oxidative phosphorylation are limiting [17, 53, 55, 63]. Our findings are consistent with a model in which peri-droplet mitochondria predominantly supply ATP and reducing equivalents for LD biogenesis, while a separate mitochondrial population dedicated to catabolism is sustained by ongoing mtDNA synthesis and biogenesis in proliferating cells. Peri-droplet mitochondria must at some point originate from the proliferating pool of mitochondria.

Previous work demonstrated that mitochondrial network morphology interacts with CPT1 activity to shape fatty acid utilization [53]. Thus, it is likely that mitochondrial specialization to serve the needs of LD metabolism is coupled to division from the proliferating mitochondrial pool.

Primary mitochondrial defects induce the upregulation of mitochondrial biogenesis via the mitochondrial UPR [10, 60]. In contrast, we found that a lipotoxic insult, specifically an imbalance between saturated and unsaturated FAs, suppressed mtDNA synthesis and mitochondrial biogenesis. FA imbalance undoubtedly disturbs membrane biogenesis, and seems to have its most dramatic effect on the architecture and function of the ER, to which the arrest of mtDNA synthesis is likely secondary [22–24]. However, we cannot rule out that FA imbalance may also alter the properties of the mitochondrial membranes, directly affecting mtDNA synthesis by altering the properties or composition of the replisome, e.g. via the recruitment of proteins like POLG2 to membrane microdomains. Since growth rate and the catabolic functions of mitochondria were barely impacted in our study, and mtDNA copy number also remained at or near its starting value, the longer term effect of fatty acid imbalance on mtDNA is hard to predict. One possibility is that the partial arrest of mtDNA synthesis heralds a more general interruption of cell growth and proliferation until lipid stress is alleviated. Another possibility is that the stress induces a repurposing of mitochondria from catabolic to anabolic functions, increasing the pool of mitochondria dedicated to supporting LD biogenesis at the expense of those that continue to undergo proliferation.

While we observed a significant decrease in mitochondrial membrane potential in cells supplied with excess FA, there were minimal effects on mitochondrial respiratory capacity or lactate efflux, indicating that cells were still able to meet their energy demands. A limitation of this study is that all experiments were conducted with cells cultured in the presence of glucose [24]. While we found that the proportion of ATP generated by OXPHOS versus glycolysis did not significantly change upon PA treatment, alternative pathways for energy production via glycolysis or peroxisomal beta-oxidation presumably remain intact in the cells. Whether our observations are specific to liver-derived cells or represent a common paradigm among eukaryotic cells remains to be explored. LDs are found widely, but can be especially abundant in the liver, e.g. in cases of non-alcoholic fatty liver disease [21, 24]. Moreover, cultured hepatocarcinoma cells are distinct from the bulk of hepatocytes in that they undergo rapid proliferation. Whether mtDNA synthesis and biogenesis are affected by saturated fatty acids in the same way in post-mitotic cells remains to be tested.

Cell culture in excess saturated FA has been linked to decreased endomembrane fluidity [64]. Thus one possibility is that the addition of monounsaturated FA restores fluidity to an extent sufficient to neutralize and traffic the free FA pool out of the ER, into LDs. In parallel, it is plausible that there exists a threshold of membrane fluidity required for the formation and stabilization of MCS between organelles. If indeed saturated lipid stress reduces ER, but not mitochondrial membrane fluidity in cultured cells, this could plausibly impact the recruitment of MCS proteins to these sites and thus MCS functions, including mtDNA synthesis.

A subset of ER-mitochondria MCS coordinate mitochondrial membrane growth and division with mtDNA synthesis, which occurs on the mtDNA copies adjacent to these sites [25, 27, 43]. Our results raise the intriguing possibility that mtDNA replisome proteins depend on the local lipid microenvironment at these contacts for their recruitment, assembly, or processivity. Recent work suggests that lipid modifications on the outer mitochondrial membrane are signals for local enrichment of receptors for the mitochondrial division dynamin, DRP1 [65]. Thus, differential composition of a lipid platform at ER-mitochondria MCS dedicated to mtDNA synthesis versus LD biogenesis could be a mechanism to specify MCS classes with dedicated functions and to coordinate the recruitment of the relevant proteins to these sites across both mitochondrial membranes.

Dysregulated mtDNA copy number is a signature of a high-fat diet, both in humans and in animal models [66, 67]. Therefore, a fuller understanding of how lipid stress impacts mtDNA metabolism provides a promising route to the identification of new mitochondrial biomarkers and therapeutic targets, whether in overnutrition, malnutrition, starvation or various forms of metabolic disease.

## Materials and Methods

### Cell culture and fatty acid treatment

Huh7 human hepatoma cells were obtained from the University of California, Berkeley Cell Culture Facility (Research Resource Identifier SCR_017924). Cells were maintained in low glucose culture media, prepared with Dulbecco’s Modified Eagle Medium (DMEM) lacking glutamine, phenol red, sodium pyruvate, and HEPES (Gibco Cat. No. A1443001), supplemented with 10% dialyzed fetal bovine serum (Gibco Cat. No A3382001), 1% L-glutamine (100X) (Gibco Cat. No. 25030081), 1% penicillin/streptomycin (Gibco Cat. No. 15140122), and 5 mM [+] D-glucose (Gibco Cat. No. A2494001). Cells were maintained at 37 °C in a humidified 5% CO_2_ chamber, on Falcon tissue culture dishes (Corning Cat. No. 353003), or 6 well, flat bottomed tissue culture plates (GenClone Cat. No. 25105). For imaging, cells were cultured in glass-bottom 35 mm dishes (Mattek) in 2 mL of complete medium. The long chain fatty acids palmitic acid (Sigma-Aldrich Cat. No. P0500) or oleic acid (Cole-Parmer Cat. No. C2110000-500C) were made up as 1 mL stock solutions at concentrations of 0.1 M and 0.3 M respectively. Fatty acid solutions were prepared by dissolving lyophilized fatty acid in dimethyl sulfoxide (DMSO). To increase solubility, the solution was warmed to 37 °C and vortexed for 1 minute or until all lyophilized material was visibly dissolved. Stock solutions of fatty acids in DMSO were stored at −20 °C. Cells were treated with a 1:1000 dilution of palmitic or oleic acid in complete culture medium for 24 h in 5% CO_2_ at 37 °C with final concentrations, as described in the main text.

### Plasmids and Transient Transfection

Cells were plated onto glass-bottom dishes 24 h prior to transient transfection. Transfections were conducted for 4-5 h in serum-free medium (Opti-MEM; Gibco Cat. No. 31985070) using Lipofectamine 2000 (Invitrogen Cat. No. 1168027). Following the 4-5 h incubation in transfection mixture, cells were treated with fatty acid, as described above, for 24 h, followed by live imaging. Plasmids and resource identifiers are reported in **Table S1**.

### General Airyscan fluorescence microscopy

All imaging was performed with an inverted 63x/1.4 NA oil objective using the laser scanning confocal Zeiss LSM 980 microscope with Airyscan 2. Images were taken with laser lines 405, 588, 561, and 639 and captured using bidirectional scanning, with a scan speed of 0.5-2 seconds.

### Live cell imaging

Live imaging was performed at 37 °C in a humidified chamber with environmental control set to 5% CO_2_. Vital dyes were used at the following final concentrations: MitoTracker Deep Red (50 nM; Invitrogen Cat. No. M22426), BODIPY 493/503 (0.67 µM; Invitrogen Cat. No. D3922), BODIPY 665/676 (50 ng/mL; Invitrogen Cat. No. B3932), BODIPY 558/568 C12 (5 µM; Thermo Scientific Cat. No. D3835), and Tetramethylrhodamine, ethyl ester (TMRE; 0.2 µM; Invitrogen Cat. No. T669). Cells were incubated in complete medium at 5% CO_2_ and 37 °C for 12-15 minutes to allow staining, with the exception of BODIPY dyes. For BODIPY 665/676 and BODIPY 558/568 C12, cells are treated with the dye for 24 h concurrent with fatty acid treatment. Reagents and pharmacologics were dissolved in DMSO and applied to cells in complete medium as indicated in **Table S2**.

### Immunofluorescence

Cells were plated onto glass-bottom 35 mm dishes (Mattek) 2-4 days before fixation, with a final confluency of 40-60%. Staining with BODIPY 493/503 was completed prior to fixation, with a final concentration of 3.8 µM with a 20-25 minute incubation.

#### 4% Paraformaldehyde fixation

Cells were fixed with 2 mL of pre-warmed (37 °C) 4% paraformaldehyde (PFA) in PBS (pH 7.4) for 20-30 minutes at room temperature, protected from light. The fixation reaction was quenched by adding 1 M glycine at a 1:100 dilution directly to the PFA in the imaging dish and incubated at room temp for 5 minutes. After quenching, samples were gently washed with PBS. Cells were then permeabilized in 2 mL of 0.1% TritionX-100 diluted in PBS for 10 minutes. Following permeabilization, cells were washed with 2 mLs of blocking buffer (1X TBS, pH 7.6, 0.1% Tween-20, 1% bovine serum albumin) for 10 minutes at room temperature, gently shaking. EdU labeling and immunolabeling followed blocking.

#### Immunolabeling

After a 10 minute wash in blocking buffer, the solution was aspirated and 2 mLs of fresh blocking buffer added to the sample. Primary antibodies were diluted 1:1000 in this solution, and incubated either at room temperature for 1 h, or at 4 °C overnight. Samples were washed with 2 mLs blocking buffer for 10 minutes, and then a 1:2000 dilution of secondary antibody in blocking buffer was added for a 1 h incubation. A final wash in blocking buffer and imaging conditions were as described above. Primary and secondary antibodies are reported in **Table S3**.

### EdU labeling of mitochondrial DNA

This protocol was adapted from a previously reported method [69]. 5-ethynyl-2’ - deoxyuridine (EdU) labeling was performed using the Click-iT™ Plus EdU Cell Proliferation Kit for Imaging, Alexa Fluor™ 647 dye (Invitrogen Cat. No. C10640), prepared according to manufacturer instructions. Click-iT® EdU buffer additive was used at a final concentration exactly ten-fold the manufacturer recommendation. We reserved 1 mL of conditioned medium from each sample. Then 1 µL of 10 mM EdU stock diluted in 0.5 mL conditioned medium was mixed thoroughly and added dropwise to cells. Cells were incubated in 5% CO_2_ at 37 °C for 30-45 minutes for pulse labeling, as described in the main text and figure legends. After the EdU pulse, the labeling medium was replaced with the 1 mL of reserved conditioned medium and samples were incubated in 5% CO_2_ at 37 °C for 10 minutes. Samples were fixed as described above. Post-permeabilization, sites of EdU incorporation were rendered fluorescent by copper click reaction. Following the click reaction, 1.5 mLs of blocking buffer was added for a brief wash, then aspirated.

### General Image quantification

2D and 3D image processing and maximum intensity projections of z-stack images were performed using Zeiss ZEN Blue software version 3.7 (Carl Zeiss). Quantification of features within Airyscan processed .czi files were analyzed as follows, using FIJI and Zeiss Arivis. Airyscan processed .czi files were converted to .sis files to be used on Zeiss Arivis Pro 4.2.1 image analysis software. Colocalization analysis of GFP-Sec61b and KDEL-mRuby in Figure 4 was performed in Fiji using the ‘coloc2’ plugin with bisection threshold regression.

### Fiji analysis of Mitochondrial area, dsDNA foci count, and EdU foci count

Airyscan processed .czi files were z-projected at maximum intensity. Trainable Weka Segmentation was used to create a classifier to detect mitochondria (Tom20 signal), mtDNA foci, and EdU foci. A region of interest mask for mitochondria was then created to discard any signal outside of segmented mitochondria. The number of mtDNA or EdU foci segmented within the mitochondrial mask were then quantified. Several images were used to create the classifiers using a macro code written in FIJI, and a custom macro was also created to quantify mitochondrial area, and mtDNA and EdU foci count.

### TMRE, Mitotracker, and BODIPY Segmentation

Machine Learning Segmentation was used for the detection of faint to bright TMRE or Mitotracker labeled mitochondria. A denoising parameter was applied to the BODIPY 493/503 channel to smooth lipid droplets, using the discrete gaussian method. Machine Learning Segmentation was applied to the BODIPY channel for classification of lipid droplets. After segmentation of lipid droplets, a watershed function was used to separate individual droplets that may be touching. A size-based object filter was used to remove objects smaller than a surface voxel area of 0.25 µm^2^ for both the TMRE and BODIPY channels. Mean and maximum signal intensity, surface voxel area, or roundness was measured from the segmented and filtered objects. Images were run as a batch analysis, and values were then exported as .xlsx files.

### POLG2-GFP foci analysis

For the segmentation of mito-BFP expressing mitochondria, a shape detection function was used for the mito-BFP channel to identify bright filaments that are 0.25 - 1 µm in length. An intensity threshold segmenter was used to classify Mito-BFP expressing mitochondria, and a size-based object filter was used to remove segmented mitochondria smaller than 0.1 µm^3^. The blob finder segmentation tool was used to classify POLG2-GFP, with a split sensitivity of 50% set to identify GFP puncta with an average diameter of 0.4 µm. A size-based object filter was used to remove segmented POLG2-GFP smaller than 0.04 µm^3^, and a signal intensity object filter was applied to remove segmented POLG2-GFP with below-threshold signal intensity. The compartmentalization function allowed for the quantification of POLG2-GFP that are inside or that intersects with segmented mitochondria. Volume and surface voxel area was measured from the segmented and filtered objects. Images were run as a batch analysis, and values were then exported as .xlsx files.

### Red C12 fatty acid lipid transfer assay

We adapted a previously reported method to monitor fatty acid trafficking in cells [68]. We defined regions of interest (ROI) delineating each respective organelle and applied an object mask filter to exclude extra-cellular signals. Images were converted to .sis files for the remainder of the analysis. In Zeiss Arivis, the blob finder segmentation tool was used to segment BODIPY 665/676 stained cells. An intensity threshold segmenter was used to classify mitochondria, and a size-based object filter was used to remove segmented mitochondria smaller than 0.1 µm^3^. Machine Learning Segmentation was used for the segmentation of the ER channel. Using the object math function, segmented mitochondria and LDs that intersected with segmented ER were subtracted from the ER. The signal intensity of the Red C12 channel was then measured within the segmented ER, mitochondria, and LDs, as well as within the entire ROI of the cell to get a whole cell measurement of Red C12 intensity. After batch analysis, values were then exported as .xlsx files. To determine the ratio of Red C12 within the organelles, named the Red C12 colocalization index, the signal intensity of Red C12 in each of the three organelles was divided by the whole cell RedC12 intensity value. The proportion of cytosolic Red C12 was determined by subtracting the sum of the organelle Red C12 ratios from 1. The density of Red C12 in each organelle was determined by dividing the Red C12 colocalization index for each organelle by organelle surface voxel area.

### Tripartite MCS Intersection analysis

Similar to the classification methods described above, ER, POLG2 foci, mitochondria, and lipid droplets were segmented and filtered. An object math feature was applied to determine all intersection points between segmented ER, mitochondria, and POLG2 (three-way intersection) per cell. After determining these three-way intersections, another object math feature was applied to identify points where the three-way intersections overlap with lipid droplet signal. Images were run as a batch analysis, and values were then exported as .xlsx to determine the ratio of three-way to four-way intersections.

### Western blotting

Cells were cultured and treated with fatty acids for 24 h in 6 well, flat bottomed tissue culture plates (GenClone Cat. No. 25105) at 40-60% confluency. Cells were trypsinized (Gibco Cat. No. 25200056), collected, and spun down at 800 x g for 2 minutes. The supernatant was aspirated, and cells were resuspended in lysis buffer (1:10 dilution of 10X TBS, 0.5% SDS dissolved in ultrapure distilled water, 0.05% TritionX-100, 5 mM EDTA, and ultrapure distilled water). Loading dye was prepared with 2x Laemmli Sample Buffer and 2-mercaptoethanol (BME), according to manufacturer instructions. A 1:2 dilution of cell lysate and loading dye was boiled for 5 minutes at 90 °C, and then spun down at 21,300 x g for 5 minutes. Samples were loaded into a SDS-PAGE gel (Invitrogen Cat. No. XP04205BOX) for electrophoresis separation. The gel was then placed in coomassie blue staining dye (Bio Rad Laboratories Cat. No. 1610786) and destained with deionized water for protein level quantification to ensure that the same amount of total protein was loaded per well. After quantifying overall protein concentration between samples, samples were prepared with loading dye (made with 4x Laemmli Sample Buffer and BME, according to manufacturer instructions) similar as before, and ran on a second SDS-PAGE gel. Proteins were transferred to a nitrocellulose gel using the iBlot NC mini-stack (Invitrogen Cat. No. IB23002). The membrane was placed in blocking buffer (see immunofluorescence protocol) and a 1:5000 dilution of primary antibody, protected from light, and incubated at 4 °C overnight. After incubation in primary antibody, the membrane was washed with blocking buffer for 10 minutes, followed by incubation in a 1:5000 dilution of secondary antibody in blocking buffer for 1 h at room temperature. The membrane was washed with blocking buffer for 10 minutes, replaced with fresh blocking buffer, and imaged on a Licor Odyssey Clx Imager.

### DNA isolation

After fatty acid treatment as described above, trypsinized cells were resuspended in 100 µL lysis buffer and kept on ice. A 1 µL aliquot of RNAse A (Thermo Scientific Cat. No. EN0531) was added to the lysed cells for a 5 minute incubation on ice, followed by the addition of 1 µL Proteinase K (Thermo Scientific Cat. No. EO0491) for an additional 5 minutes. The lysate was then diluted with 600 µL of nuclease free water and phenol-chloroform extraction was performed followed by ethanol precipitation and air-drying. DNA samples were then resuspended in 20 µLs of nuclease free water, vortexed/flicked to ensure even mixing and nano-dropped to measure the concentration of DNA. Samples were then used directly for quantitative PCR or kept at −20 °C.

### Quantitative PCR measurement of mtDNA copy number

Primer sequences are reported in **Table S4**. To prepare samples for qPCR, CutSmart buffer (New England BioLabs Cat. No. B7204S) was added to the DNA at a 1:10 dilution (e.g. 2 µL in 18 µL of DNA sample). BAMHI-HF digestion enzyme was added at 1 µL (New England BioLabs Cat. No. R3136S), and tubes were incubated at 37 °C for 40 minutes. The concentration of DNA used was 50 ng/µL, and the samples were diluted with nuclease free water as needed. qPCR primers were designed for the mitochondrial gene *rnr2* and the nuclear gene *b2m*. To prepare the qPCR reaction mix the following reagents were used: 0.1 µM of forward and reverse primers, 1:10 dilution of isolated DNA, 1:2 dilution of SsoAdvanced Universal SYBR® Green Supermix (Bio-Rad Cat. No. 1725270), and nuclease free water. As an example, one reaction included: 0.2 µL each of forward and reverse mtDNA and nuclear primers, 10 µL of SYBR green qPCR master mix, and 7.6 µL of nuclease free water. After the reaction mixture was prepared, samples were pipetted into 96 well PCR plates with 2 µL of isolated DNA and 18 µL of reaction mix, for 20 µL of solution per well. Each sample was run in triplicate. Plates were sealed, spun down 5 minutes, and the qPCR was run on a Biorad CFX Opus 96 with the following cycling conditions: denaturation for 15 seconds at 95 °C and annealing/extension for 45 seconds at 58 °C, for a total of 40 cycles. Mitochondrial genome copy number was calculated from qPCR data by averaging the Ct values of the mtDNA technical replicates and the nuclear DNA (nDNA) technical replicates. The average mtDNA Ct value was normalized to the nuclear Ct value (ΔCt) of the negative control (no treatment sample). All other ΔCt values were normalized to this value (ΔΔCt). Finally, mtDNA copy number was indicated as the fold change of the ΔΔCt.

### Seahorse XF respirometry analysis

All respirometry was conducted in a Seahorse XF96 or XFe96 Analyzer (Agilent Technologies). All experiments were conducted at 37°C and at pH 7.4. The placement of treatment groups on the XF plate was randomized across biological replicates as best as possible to avoid biased results. Respiration was measured in medium containing 8 mM glucose, 2 mM glutamine, 2 mM pyruvate, and 5 mM HEPES. Cells were plated at 2 x 10^4^ cells/well and allowed to adhere for 24 hours prior to treatment. Respiration was measured in response to oligomycin (1 µM), carbonyl cyanide-p-trifluoromethoxyphenylhydrazone (FCCP) (0.75 nM or 1.5 µM), and rotenone (0.2 µM) with antimycin A (1 µM). Calculations of respiratory parameters were made according to standard protocols [70–71] . Briefly, ATP-linked respiration was calculated by subtracting the oxygen consumption rate insensitive to rotenone and antimycin A from the measurements after injection of oligomycin. Maximal respiration was calculated by subtracting the oxygen consumption rate insensitive to rotenone and antimycin A from the maximum rate obtained after injection of FCCP. Lactate efflux rates and ATP production rates were calculated as previously described [72] .

### Cell viability and normalization

When normalizing respirometry experiments and metabolite quantification to cell number, cells were fixed immediately upon completion of the assay with 2% (w/v) formaldehyde for 20 min at room temperature and kept refrigerated between 1-14 days until assessment. On the day prior to cell counting, cells were stained with Hoechst (Thermo Fisher #33342) at 10 ng/mL overnight at room temperature. Cell counts were obtained using the Operetta High Content Imaging System (Perkin Elmer).

### Statistical analyses

All data collection, organization, and calculations were performed with Microsoft Excel. Data was plotted and statistically analyzed using Prism version 10.1.1. Plots display the mean values ± standard deviation. Statistical parameters, such as cell count and number of biological replicates (N) can be found in the figure legends. For experiments involving 2 groups, statistical significance was determined using a two-tailed unpaired t-test. For experiments involving 3 or more groups, statistical significance was determined using one-way ANOVA and Dunnett’s multiple comparisons test. For the RedC12 colocalization index in Figure 4, two-way ANOVA and Tukey’s multiple comparisons test was applied. P-values < 0.05 were considered significant.

## Supplemental material

Supplementary Figures S1-S5

Tables S1-4

## Acknowledgements

National Institutes of Health grant R00GM129456 (S.L.)

National Institutes of Health grant R35GM147218 (S.L.)

National Institutes of Health grant R35GM138003 (A.D.)

National Institutes of Health grant P30DK116074 (S.L. via Stanford Diabetes Research Center)

UC LEADS: Leadership Excellence through Advanced Degrees (B.D & A.D.)

We thank H. Jacobs and A. Arruda for close reading of this work.

## Author contributions

All data, text, and figures in the manuscript were generated by humans.

Conceptualization: C.B. and S.C.L. Methodology: C.B., S.J., B.D., T.P.W., A.B., A.D., and S.C.L. Software: C.B., S.J., and S.C.L. Validation: C.B. and S.C.L. Formal analysis: C.B., S.J., S.C.L. Investigation: C.B., S.J., B.D., H.S., A.S., T.P.W., A.B., A.D., and S.C.L. Resources: A.D., C.B., and S.C.L. Data curation: C.B., A.D., and S.C.L. Writing—original draft: C.B. and S.C.L. Writing—review and editing: C.B., S.J., and S.C.L. Visualization: C.B., S.J., H.S., and S.C.L. Supervision: S.C.L. Project administration: A.S., and S.C.L. Funding acquisition: A.D., and S.C.L.

## Abbreviations

ATP: adenosine triphosphate
CPT1: Carnitine palmitoyltransferase 1
DGAT1/2: Diglyceride acyltransferase 1 and 2
DMSO: dimethyl sulfoxide
DRP1: Dynamin-1-like protein
EdU: 5-ethynyl-2-deoxyuridine
ER: endoplasmic reticulum
ETC: electron transport chain
FA: fatty acid
FAO: fatty acid oxidation
LD: lipid droplet
MCS: membrane contact sites
mtDNA: mitochondrial DNA
PA: palmitic acid
POLG1/2: mitochondrial polymerase gamma 1 and 2
OA: oleic acid
SSBP1: single stranded binding protein 1
TMRE: tetramethyl rhodamine ester
UPR: unfolded protein response.

