## Supplementary material for "Saturated lipid stress attenuates mitochondrial genome synthesis in human cells": Table S1

**Table S1. Plasmids**

| Plasmid | Source and Reference |
|---|---|
| Mito-BFP | mito-BFP was a gift from Gia Voeltz (Addgene plasmid # 49151 ; http://n2t.net/addgene:49151 ; RRID:Addgene_49151) https://pubmed.ncbi.nlm.nih.gov/21885730/ |
| Mito-mCherry | mCherry-Mito-7 was a gift from Michael Davidson (Addgene plasmid # 55102 ; http://n2t.net/addgene:55102 ; RRID:Addgene_55102) https://www.ncbi.nlm.nih.gov/pubmed/18228502 |
| KDEL-mRuby | KDEL-mRuby was a gift from Joerg Wiedenmann https://pubmed.ncbi.nlm.nih.gov/19194514/ |
| KDEL-BFP | BFP-KDEL was a gift from Gia Voeltz (Addgene plasmid # 49150 ; http://n2t.net/addgene:49150 ; RRID:Addgene_49150) https://pubmed.ncbi.nlm.nih.gov/21885730/ |
| POLG2-GFP | POLG2-GFP was a gift from Bill Copeland https://pubmed.ncbi.nlm.nih.gov/26123486/ |
| GFP-SEC61b | GFP-SEC61b was a gift from Christine Mayr (Addgene plasmid # 121159 ; http://n2t.net/addgene:121159 ; RRID:Addgene_121159) |
| Seipin-GFP | Seipin-turboGFP was ordered from Origene. The plasmid was cloned from human mRNA NCBI accession NM_032667, human BSCL2/Seipin product catalog number RG203625 |
| SSBP1-mKate | SSBP1-mKate was custom synthesized by Origene based on NCBI accession NM_003143, human SSBP1 mRNA transcript variant 5 |
