## Supplementary material for "Saturated lipid stress attenuates mitochondrial genome synthesis in human cells": Table S2

**Table S2. Chemicals and Pharmacologics**

| Reagent/Resource | Reference Source | Identifier/Catalog No. |
|---|---|---|
| Carbonyl cyanide-p-trifluoromethoxyphenylhydrazone (FCCP)<br><br>(30 minutes, 20 μM) | Cayman Chemical Company | Cat# 15218 |
| T863 (DGAT1 inhibitor)<br><br>(48 hours, 20 μM) | MedChemExpress | Cat# HY-32219 |
| PF-06424439 (DGAT2 inhibitor)<br><br>(48 hours, 10 μM) | MedChemExpress | Cat# HY-108341A |
| Etomoxir<br><br>(2 hours, 40 μM) | Thomas Scientific | Cat# C986W52 |
| Tunicamycin<br><br>(2 hours, 12 μM) | Cell Signaling Technology | Cat# 12819S |
| ISRIB<br>(24 hours, 0.2 μM) | MedChemExpress | Cat# HY-12495A |
| Oligomycin | Sigma-Aldrich | Cat# 75351 |
| Rotenone | Sigma-Aldrich | Cat# R8875 |
| Antimycin A | Sigma-Aldrich | Cat# A8674 |
| FCCP | Sigma-Aldrich | Cat# C2920 |
| Glucose | Sigma-Aldrich | Cat# G8769 |
| Sodium Pyruvate | Sigma-Aldrich | Cat# P5280 |
| L-Glutamine | Sigma-Aldrich | Cat# G3126 |
| HEPES buffer | Sigma-Aldrich | Cat# 83264 |
| DMEM Powder | Sigma-Aldrich | Cat# 5030 |

| | | |
|---|---|---|
| Sodium Chloride | Sigma-Aldrich | Cat# S7653 |
| Phenol Red | Sigma-Aldrich | Cat# P0290 |
| Water suitable for cell culture | Sigma-Aldrich | Cat# W3500 |
| Dulbecco's phosphate-buffered saline (DPBS) | Fisher Scientific | Cat# 14-190-250 |
| 16% Paraformaldehyde | Fisher Scientific | Cat# 50-980-487 |
| Hoescht 33342 | Fisher Scientific | Cat# H3570 |
