## Supplementary material for "Saturated lipid stress attenuates mitochondrial genome synthesis in human cells": Table S3

**Table S3. Primary and Secondary antibodies**

| Description | Source | Catalog No. |
| --- | --- | --- |
| TOM20 Polyclonal Antibody | Proteintech | 11802-1-AP |
| Anti-dsDNA antibody [35I9 DNA] - BSA and Azide free | abcam | ab27156 |
| Calreticulin Polyclonal Antibody | Invitrogen | PA5-34786 |
| GRP78 Polyclonal Antibody | Invitrogen | PA1-014A |
| CHOP Polyclonal Antibody | Proteintech | 15204-1-AP |
| XBP-1s (E9V3E) Rabbit mAb | Cell Signaling Technology | 40435S |
| β-Actin (13E5) Rabbit mAb | Cell Signaling Technology | 4970 |
| Goat anti-Mouse IgG (H+L) Cross-Adsorbed Secondary Antibody, Alexa Fluor™ 405 | Invitrogen | A31553 |
| Donkey anti-Rabbit IgG (H+L) Highly Cross-Adsorbed Secondary Antibody, Alexa Fluor™ Plus 488 | Invitrogen | A32790 |
| Donkey anti-Rabbit IgG (H+L) Highly Cross-Adsorbed Secondary Antibody, Alexa Fluor™ Plus 680 | Invitrogen | A32802 |
| Goat anti-Mouse IgG (H+L) Highly Cross-Adsorbed Secondary Antibody, Alexa Fluor™ Plus 800 | Invitrogen | A32730 |
