## Supplementary material for "Saturated lipid stress attenuates mitochondrial genome synthesis in human cells": Table S4

**Table S4. Primers for quantitative PCR of mtDNA copy number**

| Gene | Forward Primer | Reverse Primer |
|---|---|---|
| NM_004048.4 Homo sapiens beta-2-microglobulin (B2M) mRNA | TGCTGTCTCCATGTTTGATGTATCT | TCTCTGCTCCCCACCTCTAAGT |
| NC_012920.1:1671-3229 Homo sapiens mt-Rnr2 | AGACTTCACCAGTCAAAGCGA | ACATCGAGGTCGTAAACCCT |
