## Supplemental Figures S1-S5 for "Saturated lipid stress attenuates mitochondrial genome synthesis in human cells"

### Figure S1

A

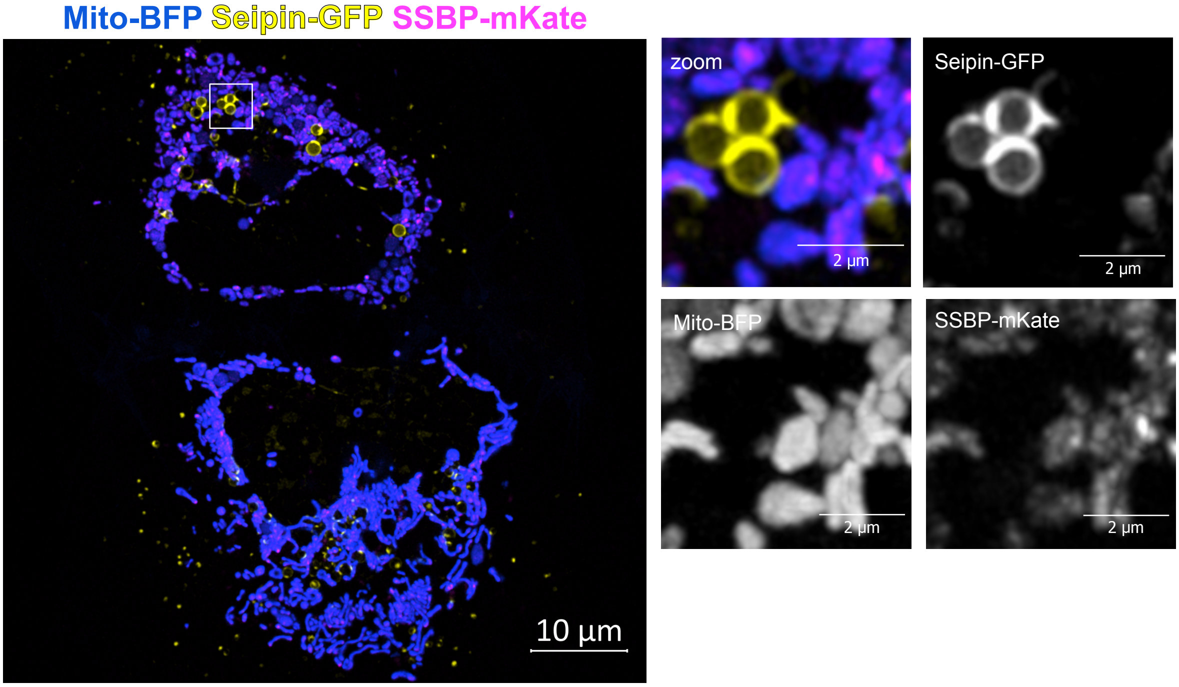
Mito-BFP Seipin-GFP SSBP-mKate
10 µm
zoom
2 µm
Seipin-GFP
2 µm
Mito-BFP
2 µm
SSBP-mKate
2 µm

**Figure S1. SSBP1 localization relative to lipid droplet biogenesis marked by Seipin-GFP. (A)** Huh-7 cell transiently transfected with mito-BFP (blue), and SSBP1-mKate (magenta), and seipin-GFP (yellow) imaged by laser scanning confocal microscopy. Scale bars: 10 μm (whole cell). N = 26 cells were imaged from 2 biological replicates.

### Figure S2

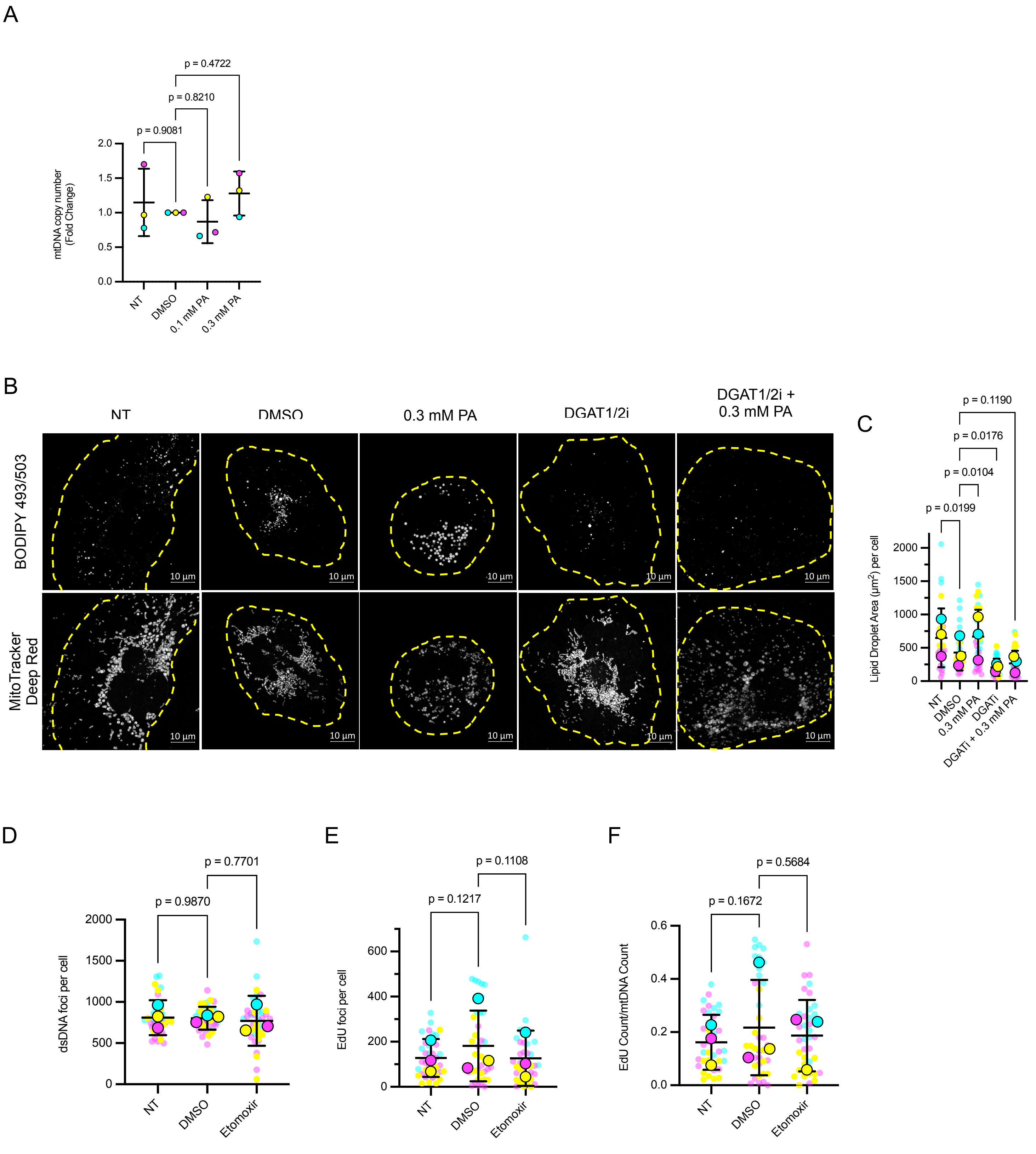

A
p = 0.4722
p = 0.8210
p = 0.9081
mtDNA copy number (Fold Change)
NT
DMSO
0.1 mM PA
0.3 mM PA
B
NT
DMSO
0.3 mM PA
DGAT1/2i
DGAT1/2i + 0.3 mM PA
BODIPY 493/503
MitoTracker Deep Red
10 µm
C
p = 0.1190
p = 0.0176
p = 0.0104
p = 0.0199
Lipid Droplet Area (µm2) per cell
DGATi
DGATi + 0.3 mM PA
D
p = 0.7701
p = 0.9870
dsDNA foci per cell
Etomoxir
E
p = 0.1108
p = 0.1217
EdU foci per cell
F
p = 0.5684
p = 0.1672
EdU Count/mtDNA Count

**Figure S2. Inhibition of fatty acid routing to LDs, but not routing to mitochondria, decreases EdU labeling of mitochondrial DNA.**
**(A)** Fold change in mtDNA measured by quantitative PCR normalized to nuclear DNA and DMSO vehicle.
**(B)** Representative fluorescence micrographs of Huh-7 cells stained with BODIPY 493/503 and MitoTracker Deep Red and treated as in B-D.
**(C)** Lipid droplet area ($\mu m^2$) per cell as segmented from Bodipy 493/503 fluorescence in DMSO vehicle, PA, DGATi, or PA+DGATi. For B-C, N = 32 cells (NT), 34 cells (DMSO), 36 cells (0.3 mM PA), 36 cells (DGATi), and 36 cells (DGATi + 0.3 mM PA), from 3 biological replicates.
**(D)** Quantification of the number of dsDNA foci per cell DMSO versus etomoxir.
**(E)** Quantification of the number of EdU foci per cell DMSO versus etomoxir.
**(F)** Calculation of the proportion of mtDNA nucleoids engaged in DNA synthesis on a per-cell basis. For D-F, N = 40 cells (NT), 40 cells (DMSO), 38 cells (Etomoxir) cells from 3 biological replicates. Statistical analysis by one-way ANOVA (Dunnett’s multiple comparisons test).

### Figure S3

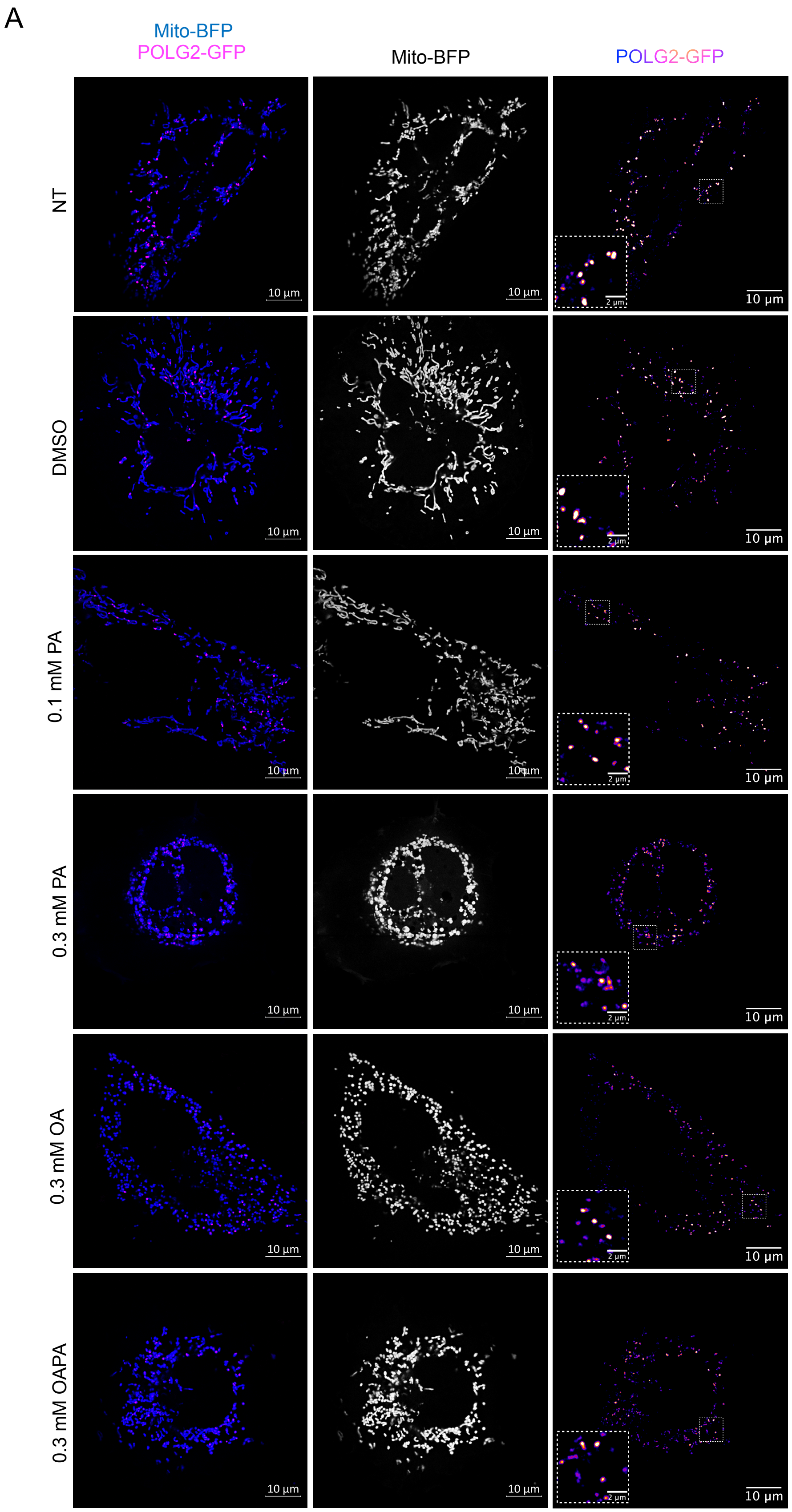
A
Mito-BFP
POLG2-GFP
Mito-BFP
POLG2-GFP
NT
DMSO
0.1 mM PA
0.3 mM PA
0.3 mM OA
0.3 mM OAPA
10 µm
2 µm

**Figure S3. POLG2-GFP foci in cells treated with PA, OA, or both. (A)** Huh-7 cells transiently transfected with mito-BFP and POLG2-GFP then treated for 24 hours: Non-treated (NT), DMSO (vehicle), 0.1 mM PA, 0.3 mM PA, 0.3 mM OA, and 0.3 mM OAPA. Scale bars, 10 μm. N = 35 (NT), 30 (DMSO), 30 (0.1 mM PA), 30 (0.3 mM PA), 33 (0.3 mM OA), 30 (0.3 mM OAPA) cells were imaged from 3 biological replicates. The mito-BFP channel is shown in greyscale and POLG2-GFP channel with a ‘fire’ LUT.

### Figure S4

A

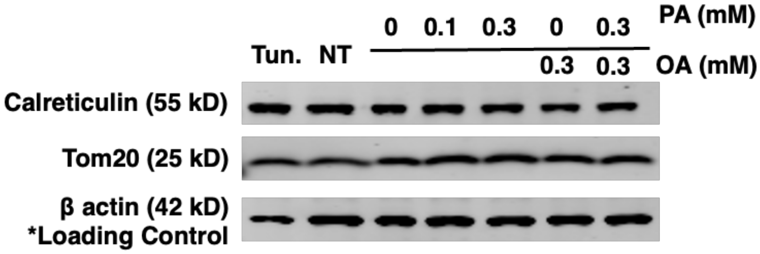


B

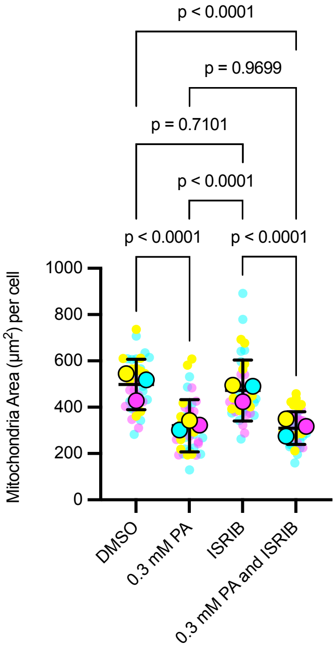


C

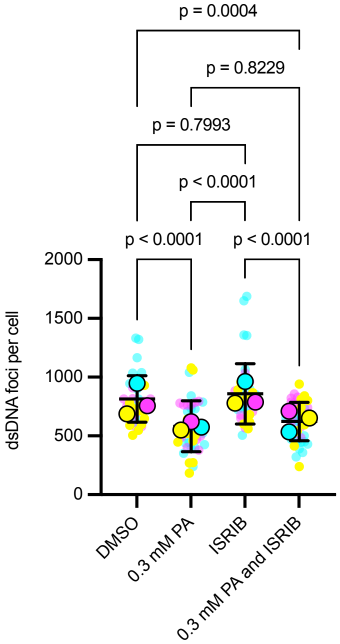


D

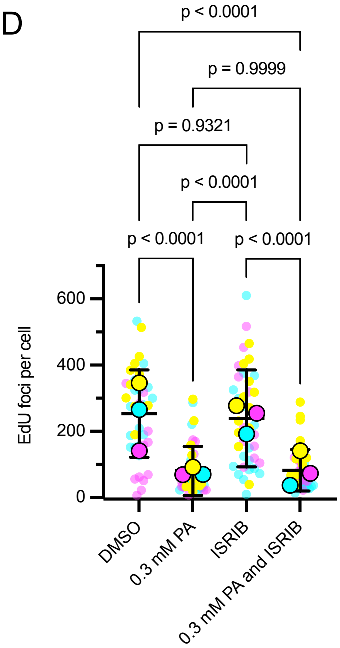


E

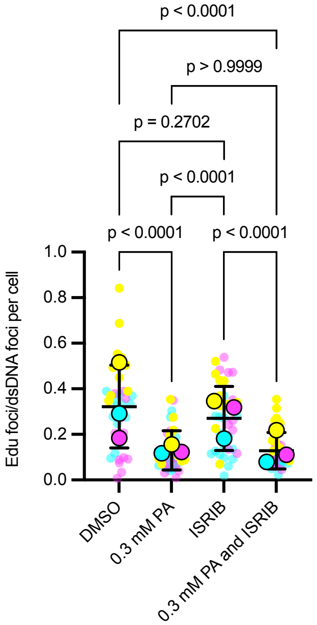

**Figure S4. ISRIB does not modify the effect of saturated fatty acids on mtDNA synthesis.**
**(A)** Immunoblot of Calreticulin, Tom20, and β-actin in Huh-7 cells treated for 24 hours as follows: Tunicamycin positive control (Tun.), No treatment negative control (NT), DMSO (vehicle), 0.1 mM PA, 0.3 mM PA, 0.3 mM OA, and 0.3 OA+PA.
**(B)** Quantification of mitochondrial area ($\mu m^2$) in ISRIB- and PA-treated cells.
**(C)** Quantification of the number of dsDNA foci per cell.
**(D)** Quantification of the number of EdU foci per cell.
**(E)** Quantification of the number of EdU foci per mitochondrion divided by the number dsDNA foci per mitochondrion.
For B-E, N = 39 (DMSO), 41 (0.3 mM PA) cells, 40 (0.2 μM ISRIB), 42 (0.2 μM ISRIB and 0.3 mM PA combined) from 3 biological replicates. Statistical analysis by one-way ANOVA (Dunnett's multiple comparisons test).

### Figure S5

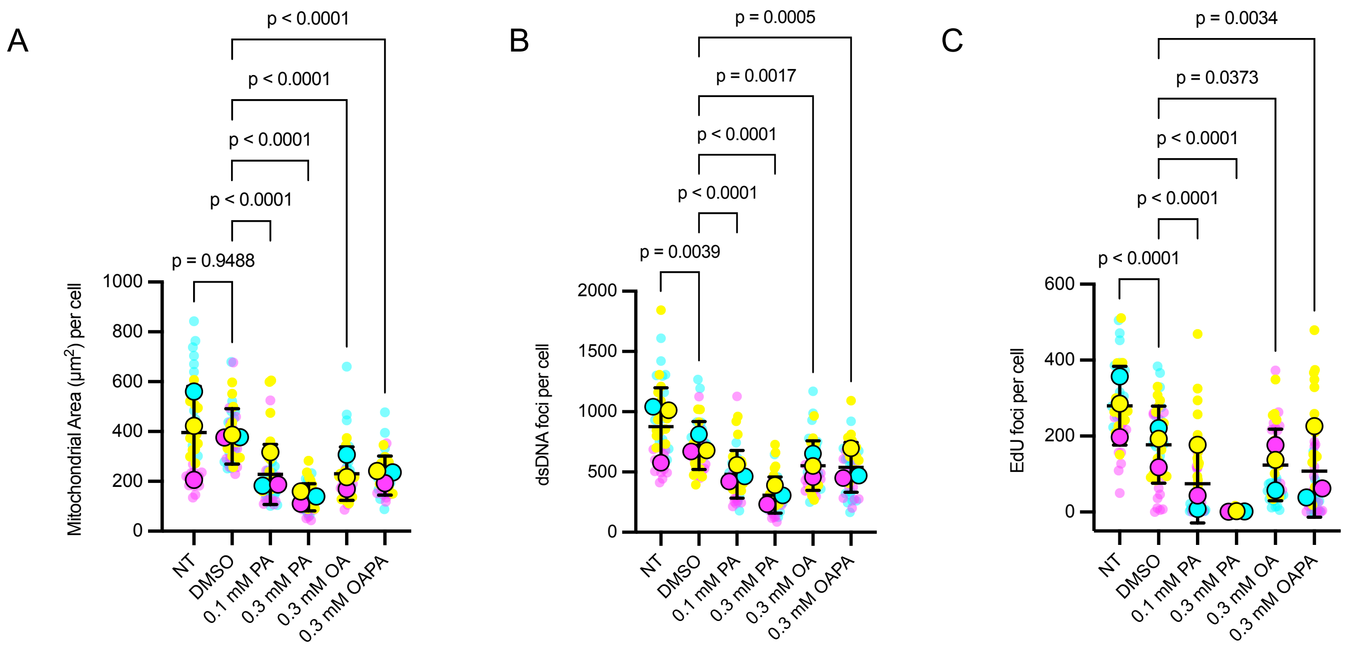
A
B
C
p < 0.0001
p < 0.0001
p < 0.0001
p < 0.0001
p = 0.9488
Mitochondrial Area (µm²) per cell
p = 0.0005
p = 0.0017
p < 0.0001
p < 0.0001
p = 0.0039
dsDNA foci per cell
p = 0.0034
p = 0.0373
p < 0.0001
p < 0.0001
p < 0.0001
EdU foci per cell
NT
DMSO
0.1 mM PA
0.3 mM PA
0.3 mM OA
0.3 mM OAPA

**Figure S5. Mitochondrial parameters in cells treated with oleic acid.**
**(A)** Quantification of mitochondrial area ($\mu m^2$) in OA- and PA-treated cells.
**(B)** Quantification of mitochondrial dsDNA foci per cell.
**(C)** Quantification of mitochondrial EdU-647 foci per cell.
Same dataset as shown in Figure 5C-E, without normalization to mitochondrial network area. For (C-E), N = 50 cells (NT), 55 cells (DMSO), 61 cells (0.1 mM PA), 52 cells (0.3 mM PA), 52 cells (0.3 mM OA), 49 cells (0.3 mM OA+PA), from 3 biological replicates.
